# Volume regulation in adhered cells: roles of surface tension and cell swelling

**DOI:** 10.1101/2022.08.24.505072

**Authors:** Ram M. Adar, Amit Singh Vishen, Jean-François Joanny, Pierre Sens, Samuel A. Safran

## Abstract

The volume of adhered cells has been shown experimentally to decrease during spreading. This effect can be understood from the pump-leak model, which we have extended to include mechano-sensitive ion transporters. We identify a novel effect that has important consequences on cellular volume loss; cells that are swollen due to a modulation of ion transport rates are more susceptible to volume loss in response to a tension increase. This effect explains in a plausible manner the discrepancies between three recent, independent experiments on adhered cells, between which both the magnitude of the volume change and its dynamics varied substantially. We suggest that starved and synchronized cells in two of the experiments were in a swollen state and, consequently, exhibited a large volume loss at steady state. Non-swollen cells, for which there is a very small steady-state volume decrease, are still predicted to transiently lose volume during spreading due to a relaxing viscoelastic tension that is large compared with the steady-state tension. We elucidate the roles of cell swelling and surface tension in cellular volume regulation and discuss their possible microscopic origins.

## I. INTRODUCTION

The volume of living cells is regulated via numerous mechanisms and across different timescales, and contributes to cellular homeostasis [1–3]. Volume regulation occurs at the single-cell level. At short timescales of seconds, it is dominated by water permeation through the cell membrane and aquaporin channels. At intermediate timescales of minutes, volume is regulated mostly via ion transport through ion channels and active pumps. The transport allows the cell volume to adapt to transient changes of its environment. *In vitro* experiments allow for control of the physical surroundings of the cell over longer time scales of tens of minutes. This can result in volume changes induced, for example, by changes of the macromolecular concentration or ionic strength of the buffer.

A benchmark theoretical framework for describing cellular volume regulation is the “pump and leak” model [4, 5]. The cellular volume is determined by its mechanical equilibrium with its environment. Ions diffuse passively (termed “leak”) through channels and are transported actively by energy consuming pumps. At steady state, there is zero net flux of ions into and out of the cell. In the absence of active pumps and of macromolecules in the cellular environment, the ionic chemical equilibrium and the confinement of larger molecules (e.g., amino acids and proteins) to the cell, would lead in theory to unbounded cell volumes [5] and, in practice, to cell lysis.

Unlike passive systems in thermodynamic equilibrium, the osmotic pressure of living cells may depend on surface tension even when they are well spread and relatively flat. This is due to mechano-sensitive ion transporters that change the ion flux [6, 7] in response to surface tension and, consequently, the steady-state ionic concentrations. Heuristically, the tension can be thought to “open” closed ion transporters and increase their effective conductance. This modifies the cellular osmotic pressure and results in volume change. The dependence of cell volume on surface tension implies a possible dependence on cell shape, which can be examined during cell spreading.

Three recent, independent experiments on different cell types show that the volume of adhered cells decreases during spreading [8–10]. These studies differ in the magnitude of the effect and its dynamics. The volume decrease measured by Guo et al. [8] and Xie et al. [9] is a steady-state effect that persists throughout the experimental time scale of the order of an hour. It is independent of how the contact area is changed (*e.g*., by dynamic spreading or by patterning of substrate adhesion molecules). The volume loss in both experiments is of the order of 50%. On the other hand, the volume decrease measured by Venkova et al. [10] is transient and small, of the order of 5 – 10% for untreated cells and up to 20% under pharmacological treatments that increase the spreading rate. The initial cellular volume is recovered within about an hour. These substantial qualitative differences are not yet understood.

All three experiments emphasize the importance of cell activity for the volume decrease. The effect diminishes when activity is suppressed (e.g., via ATP depletion). Both sets of experiments invoked the pump-leak model adapted to include changes in the transport rates of ions as the cell area changes during spreading. The steady-state effect of Refs. [8, 9] was explained in Ref. [11], where the steady-state ionic chemical potential, which is determined by the balance of ion influx and efflux in the presence of active pumping, was related to the cell area. The transient effect of Ref. [10] was explained by the response of mechano-sensitive ion channels to a transient tension increase with respect to a homeostatic tension. This tension was assumed to relax via several mechanisms (*e.g*., cortical actin turnover [12]).

Here we present a unified theory for cell volume changes during spreading, induced by active ion transport and its modulation by cell tension. This allows us to explain all three experiments in an integrated manner. We propose that much of the difference between the experimental results can be attributed to different degrees of initial cell swelling due to variations in ion transport rates. We infer from the experimental measurements that the cells in Refs. [8, 9] had lower densities compared with those in Ref. [10]. Importantly, we show that such swollen cells are more susceptible to volume loss in response to a tension increase, because of the inherent non-linearity of the pump-leak model. This nonlinearity effectively amplifies the effects of tension and area changes on the volume of swollen cells. A small increase in the steady-state tension, an order of magnitude smaller than the transient tension increase during dynamic spreading, can explain the large volume loss measured in the swollen cells of Refs. [8, 9]. The same tension has little effect on non-swollen cells, like those measured in Ref. [10].

## II. METHODS

The theoretical methods of the pump-leak model and viscoelasticity were used in order to predict the cellular pressure as a function of the spreading area and spreading rate, as is detailed below.

### A. Pump leak model

Our model of the experiments considers a cell of volume *V*, which is adhered to a substrate with contact area *A_c_* and whose apical area is in contact with a buffer. The total cell area is *A*. We rely on a minimal description of the cell, consisting of water, trapped molecules that are confined to the cell (such as proteins, phosphates, and amino-acids) and ions that exchange with the buffer actively through pumps and passively through channels (“pump and leak”). This coarse-grained picture suffices to derive an expression for the osmotic pressure in the cell and, ultimately, for the cell volume, as a function of the cell area and spreading rate.

Some of the trapped molecules are mobile and contribute to the osmotic pressure. We denote their number as *N*. In order to account for excluded volume, we define an effective concentration of mobile, trapped molecules in their available volume, *n_p_* = *N* / (*V – V_d_*). Here, *V_d_* is the dried cell volume (volume of the dry mass). This is the cell volume in the limit of infinite compression. The trapped molecules are negatively charged [6] on average and attract neutralizing cations that are relatively localized [13, 14]. The cell also includes additional cations and anions of equal concentrations, which we term “additional salt”. We coarse-grain the neutralizing cations together with the trapped osmolytes and treat them as a single neutral species. We similarly treat the additional salt as a single neutral species. The charged case can be treated using the Donnan framework [15, 16] and leads to similar results (see Appendix A).

The ions exchange with the buffer, where their concentration is *n_b_*. The steady-state ion concentration in the cell n_c_ is determined by the global balance of fluxes into and out of the cell through ion channels (diffusion) and active pumps. Active pumping results in a non-equilibrium chemical potential difference between the cell and the buffer *k*_B_*Tδ*, such that *n_c_* = *n_b_* exp(–*δ*). The chemical potential difference *δ* is the ratio between the pumping rate per unit area and ion conductance [11]. In a more detailed, ion-specific description, *δ* can be related to the membrane potential (and see Sec. A of Supplementary Material).

The cells are in mechanical equilibrium with the buffer. This is described, to a good approximation, by osmotic pressure equality, because the Laplace pressure due to the cellular surface tension is several orders of magnitude smaller [8, 11] than the osmotic pressure exerted by the ions and trapped molecules. The osmotic pressure in the cell *P_c_* is obtained by adding the concentrations of the trapped molecules and salt, *P_c_* = *k*_B_*T*[*n_p_* + *n_b_* exp (–*δ*)]. The osmotic pressure in the buffer is *P_b_* = *k*_B_*Tn_b_*. Because the number of trapped molecules is fixed, *n_p_* changes only due to volume variations. The pressure equality *P_c_* = *P_b_*, therefore, relates the cell volume to *δ* via

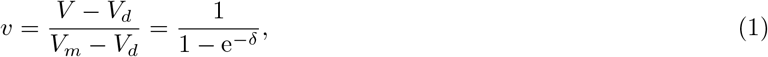

where we have defined the volume *V_m_* = *V_d_* + *N/n_b_*, for which *v* = 1. This is the minimal possible cell volume in isotonic conditions, obtained in the limit of *δ* → ∞, corresponding to infinite ion pumping out of the cell. The buffer pressure is balanced solely by the trapped molecules in this limit.

### B. Mechano-sensitivity

Some ion channels are mechano-sensitive, i.e. the ion flux across them increases in response to an increase in surface tension [6, 17]. Generally, ion pumps can be mechano-sensitive as well. Since *δ* depends on active and passive transport, it depends on the surface tension as well. The volume decrease during spreading, throughout which the tension increases, indicates that *δ* is an increasing function of the tension. This can be justified by a more detailed, ion-specific treatment [10] (and see Sec. B of Supplementary Material).

We consider small variations in *δ* due to an increase of surface tension with respect to a reference tension in suspension, *γ*_0_. We expand to linear order,

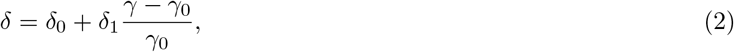

where *δ*_0_ determines the cell volume in the reference state and *δ*_1_ is a linear coefficient due to changes in mechano-sensitive transporters as the tension is increased from *γ*_0_. The general dependence of *δ* on the surface tension can be obtained from a more detailed model of mechano-sensitive ion transporters (see Sec. B of Supplementary Material).

The surface tension originates from the cell membrane and cortical layer and involves several physical mechanisms, such as actomyosin contraction and membrane fluctuations. Tension in both the membrane and cortex can relax via turnover with the cytoplasm or other cellular compartments, and the membrane-cortex coupling may similarly relax [12]. The tension can be written as *γ* = *γ_ss_* + *γ_t_*, where *γ_ss_* is the steady-state tension, and *γ_t_* is a transient tension that vanishes at steady state.

The tension at steady-state can be related, for example, to the exchange of lipids between the cell membrane and cellular organelles as part of endo- and exocytosis [18]. It can vary due to different endo- and exocytosis rates and different values of the surface tension in lipid reservoirs. The tension *γ_t_* is related to the transient elastic response of the cortex and membrane as well as to the long-time viscous stresses due to flow during spreading.

As a simple model for the tension, we consider a linear expansion around the reference state that is given by the tension *γ*_0_ and the total area *A*_0_. The strain is defined as *ϵ* = *A/A*_0_ – 1. We expand the steady-state tension in the cell area, *γ_ss_* = *γ*_0_ + *k_ss_ϵ*, where *k_ss_* is a steady-state elastic modulus. The transient tension is described by a Maxwell model, (1 + *τ∂_t_*) *γ_t_* = *k_t_τ∂_t_ϵ*, where *τ* is a viscoelastic relaxation time and *k_t_* is the transient elastic modulus. The viscosity is *η* = *k_t_τ* [19]. Together, this yields a dynamical equation for the total tension *γ* = *γ_ss_* + *γ_t_*

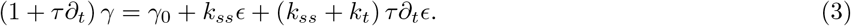

Equation (3) predicts the time-dependent tension *γ*(*t*) as a function of the history of strains *ϵ*(*t′*) for *t′* < *t*. It describes the rheology of a standard linear solid (Zener) model [20]. The system behaves as an elastic solid at short times (tension proportional to the strain) with an elastic modulus *k_ss_* + *k_t_*, then behaves as a viscoelastic solid (with tension proportional to both the strain and strain rate), and is solid again at long times with a smaller modulus *k_ss_*. An additional long-time relaxation of cellular stresses [21] is sometime considered. As the volume loss in Refs. [8, 9] persisted throughout the experimental time scale, we are not concerned with this effect and retain our simpler rheological model.

Our theory relates the cell volume to ion transport via the pump-leak model of Eq. (1). Ion transport is related to surface-tension by mechano-sensitivity, as described in Eq. (2). Finally, surface-tension is related to the cellular surface area and spreading rate according to the rheological model of Eq. (3). The combination of these equations relates the cell volume to its area and spreading rate.

The theories of Refs. [10, 11] are obtained as two limiting cases of the current theory. At steady-state, the cell volume depends only on its area and is independent of the dynamics, *γ_t_* =0. For well-spread cells, the total area is proportional to the contact area. In this limit, our theory is similar to that of Ref. [11]. The theory of Ref. [10] is obtained for dynamically spreading cells (*γ_t_* > 0) with a fixed steady-state tension *γ_ss_* = *γ*_0_ and no residual tension (*k_ss_* = 0). Only small *δ* variations were considered in Ref. [10], such that the right-hand-side of Eq. (1) can be linearized.

## III. RESULTS

### A. Swollen cells

Equations (2)-(3) are linear expansions around the reference steady-state, described by *A* = *A*_0_, *γ* = *γ*_0_ and *δ* = *δ*_0_. We consider the reference state to be that of a cell in suspension with a volume *V_s_*. This is a much less constrained situation for a cell, compared with its spreading on a substrate. In this case, Eq. (1) relates the normalized volume in suspension *v_s_* = (*V_s_* – *V_d_*) / (*V_m_* – *V_d_*) to *δ*_0_ (see Fig. 1). It shows that *v_s_* increases non-linearly, as *δ*_0_ decreases.

**FIG. 1:**
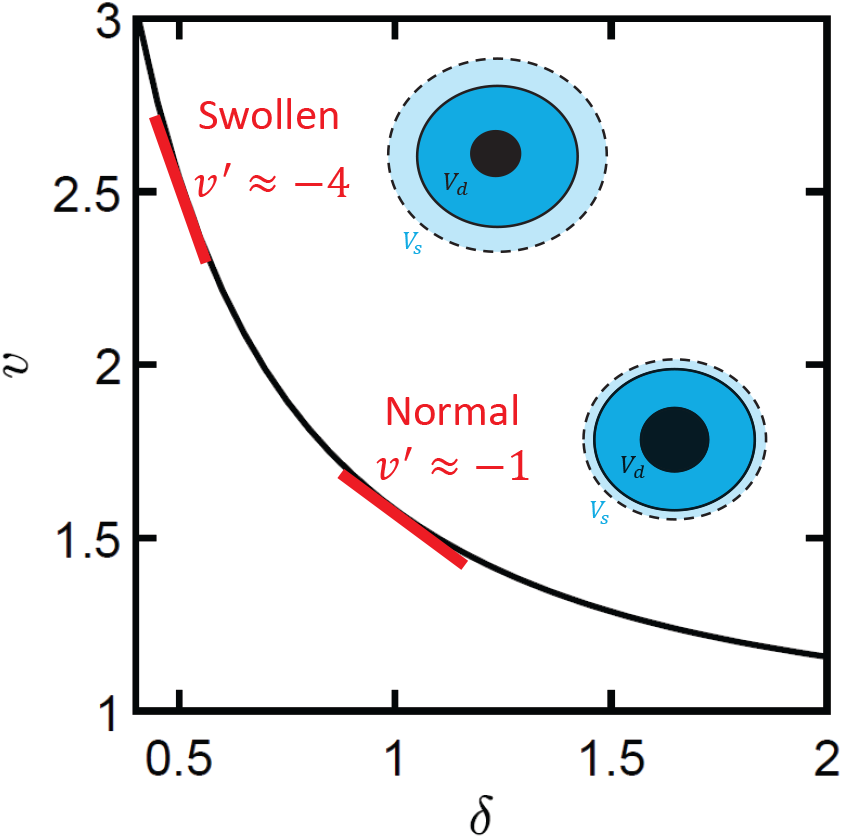
(Color online) dimensionless volume *v* as a function of the dimensionless chemical potential difference *δ*. The slope of the function is marked in red for two typical values: one of normal cells with *v* ≈ 1.7 and one of swollen cells with *v* ≈ 2.5. Swollen cells have smaller fractions of dried volume in suspension (smaller *V_d_/V_B_* values).

This nonlinear dependence originates from the exponential relation between concentration and chemical potential. The non-linearity is most evident from the limits of zero and infinitely rapid pumping (*δ*_0_ = 0 and *δ*_0_ → ∞, respectively). The cellular volume diverges for *δ*_0_ = 0, because of the pressure exerted by the trapped osmolytes, but it reaches the finite volume *V_m_* for *δ*_0_ → ∞. This non-linearity is inherent to the pump-leak model and has important implications for the volume loss during spreading. Eqs. (1) and (2) show that the effect of a tension increase with respect to the reference state is amplified by a factor exp (–*δ*_0_). This amplification is largest for small values of *δ*_0_ (large values of the normalized volume in suspension, *v_s_*) which we identify with the situation of swollen cells. Swollen cells with larger *v_s_* are thus more susceptible to volume loss. This is illustrated in Fig. 1.

The value of *v_s_* depends on two volume ratios: *V_m_/V_d_* and *V_s_/V_d_*. The former is a function of the buffer ion concentration, *n_b_*, the dried volume *V_d_*, and the number of mobile, trapped molecules *N*. It is assumed here not to vary much between different cells in the same buffer with fixed ionic concentrations (see Discussion). Therefore, we consider *v_s_* to depend mostly on *V_s_/V_d_* and swollen cells are those that have a large volume in suspension compared to their dried volume, due to smaller *δ*_0_ values. This means that even in their reference state in suspension, swollen cells have modified pumping rates and ion-channel conductance, compared to unswollen cells. Within an ion-specific model, such swollen cells have less negative membrane potentials (see Sec. A of Supplementary Material).

The higher susceptibility of swollen cells to volume loss can explain the differences in the volume decrease measured by Guo et al. and Xie et al. [8, 9], as opposed to that measured by Venkova et al [10]. Separate measurements of the same cell types indicate that the cells used in Refs. [8, 9] were larger than those used in [10]. The measured volumes of 3T3 fibroblasts and HeLa Kyoto cells were of order 2000 *μ*m^3^ in [10] and 4000 *μ*m^3^ in [8, 9]. It is plausible that these cells were not just large (implying a larger number of trapped molecules *N*), but rather swollen with smaller *V_d_/V_s_* values. Venkova et al. inferred *V_d_/V_s_* ≈ 0.3 from osmotic shock experiments, using Ponder’s relation [22]. A similar value was found in other experiments [23]. This ratio was not measured in Refs. [8, 9]. A possible reason for a modulation of ion transport rates and consequent cell swelling is the synchronization protocols used in Refs. [8, 9]. They may have affected the composition of ion channels and pumps in the cell and their performance, leading to a smaller value of *δ*_0_. We test this hypothesis further below, by comparing our theory with the experimental data.

### B. Comparison with experiments

We compare our predictions for the cell volume as a function of its area and its spreading rate to the experimental findings of Refs. [8–10]. Our theory depends on four dimensionless parameters: *V_m_/V_d_, δ*_0_, *δ*_1_ *k_ss_/γ*_0_, *k_t_/k_ss_*, and the relaxation time *τ*. The first three can be found at steady state and the final two only during dynamic spreading. Note that *δ*_0_ is directly related to the ratio *V_s_/V_d_*, as was explained above.

We fit simultaneously the two steady-state data sets of volume vs. area of Refs. [8, 9], while treating *δ*_0_ as a fit parameter and setting *k_t_/k_ss_,τ* = 0 (resulting in no transient tension, = 0). We then fit the dynamic spreading data of Ref. [10], while setting *δ*_0_ according to the measured ratio *V_d_/V_s_* = 0.3 and treating *k_t_/k_ss_* and *τ* as fit parameters. We assume the same value of *V_m_/V_d_* for all three experiments.

The fit requires the value of either *V_d_* or *V_m_* in each experiment. Guo et al. performed osmotic compression experiments on A7 cells and measured a minimal volume of about 2100 *μ*m^3^. We interpret it as the dried volume *V_d_. V_m_* was not measured. Xie et al. demonstrated that as the spread area increased, the cellular volume decreased and approached a minimal value of about 1500 *μ*m^3^. Within our theory, the minimal volume during spreading is obtained for the maximal *δ*. For simplicity, rather than introducing another model parameter, we interpret this volume as *V_m_* that corresponds to *δ* ≫ 1. The volume *V_d_* was not measured. Venkova et al. measured a volume in suspension *V_s_* ≈ 1900 *μ*m^3^ and *V_d_/V_s_* = 0.3, while *V_m_* was not measured.

As part of the fits, the contact area *A_c_* and cell volume V are related to the total area *A* using the spherical cap approximation. The reference area *A*_0_ is the surface area of the (spherical) cell in suspension. Finally, we note that the cells of Ref. [10] grow during their dynamic spreading at a rate of *g* ≈ 4%/h. Assuming that the dried volume grows at the same rate, we account for growth by multiplying *V_d_* and *V_m_* in Eq. (1) by 1 + *gt*, where *t* is the time.

Our theory fits well with the experimental data, as is evident from Fig. 2. The best fit to the data of Refs. [8, 9] is plotted for *V_m_/V_d_* = 2.5, *δ*_0_ = 0.54 and *δ*_1_*k_ss_/γ*_0_ = 0.7. The best fit to the data of Ref. [10] is plotted for *V_m_/V_d_* = 2.5, *δ*_0_ = 1, and *δ*_1_*k_ss_/γ*_0_ = 0.35, as well as *k_t_/k_ss_* = 63, and *τ* = 0.8 minutes. As part of our fits to the data of Ref. [10], we distinguish between the steady-state tension contribution to the volume loss (dashed black curve) and the full tension, including the transient tension (solid red curve).

**FIG. 2:**
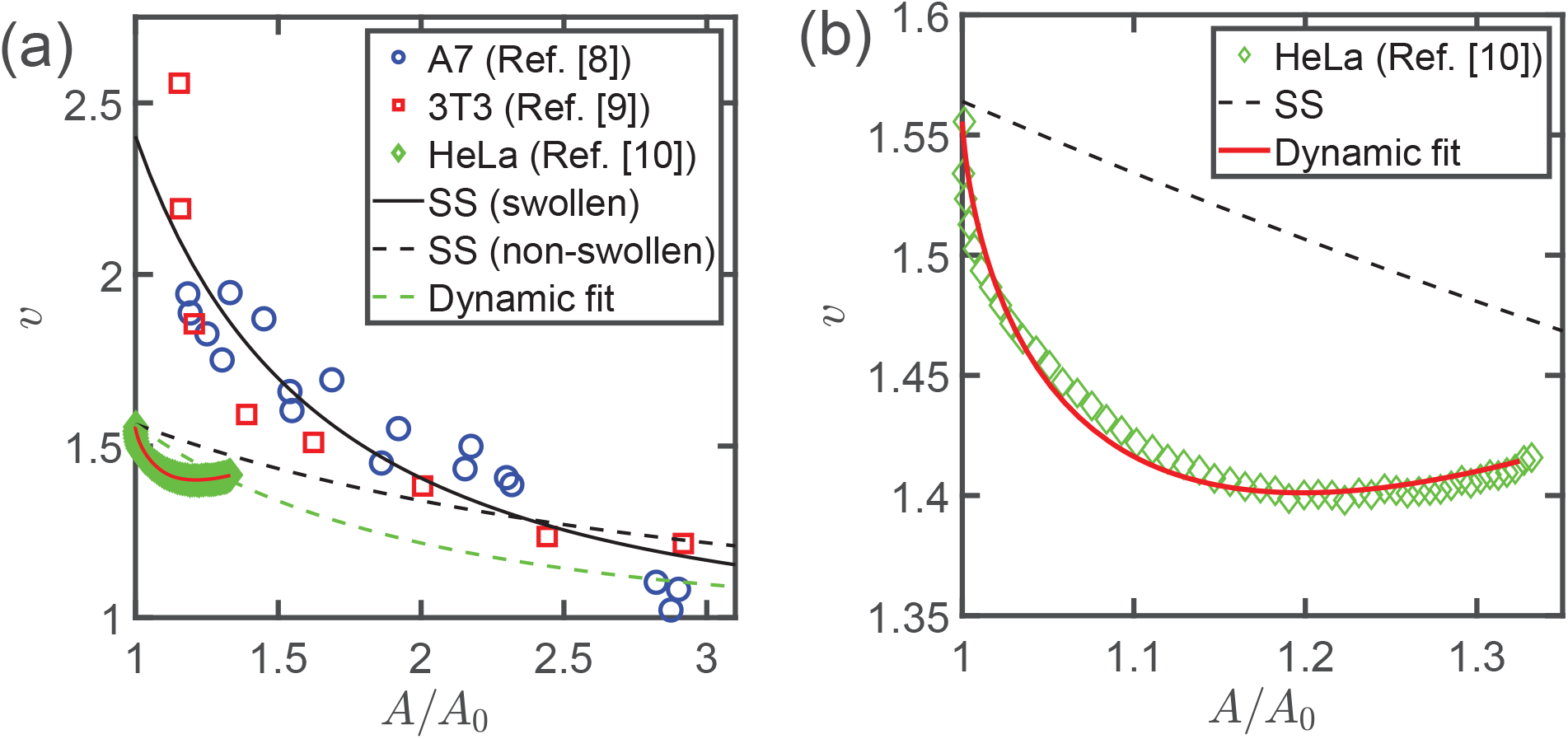
(Color online). Comparison of the theory with experimental data of cell volume as a function of cell area. The steady-state fits to Refs. [8, 9] (solid black) is plotted for *V_m_/V_d_* = 2.5 (common to all curves), *δ*_0_ = 0.54, *δ*_1_*k_ss_/γ*_0_ = 0.7, and *k_t_* = *τ* = 0. The fit to Ref. [10] (solid red) is plotted for *δ*_0_ = 1, *δ*_1_*k_ss_/γ*_0_ = 0.35, *k_t_/k_ss_* = 63, and *τ* = 0.8 minutes, according to the dynamic spreading data. The steady-state contribution to this plot (dashed black) is plotted for *δ*_0_ = 1, *δ*_1_*k_ss_/γ*_0_ = 0.35, and *k_t_* = *τ* = 0. (b) Zoom-in to the region of small volumes and areas, corresponding to the data of Ref. [10].

The value *δ*_0_ = 0.54 corresponds to *V_d_/V_s_* = 0.22, demonstrating that the cells are indeed swollen, compared to those of Ref. [10]. We note that even smaller values of *V_d_/V_s_* ≈ 0.1 were inferred recently [24] in HeLa cells from optical diffraction tomography. The best-fitted values of *δ*_1_*k_ss_/γ*_0_ are comparable for both sets of experiments. The main difference between the fit to Refs. [8, 9] (solid black line) and the steady-state contribution to the fit to Ref. [10] (dashed black curve) results from the different values of *δ*_0_, rather than those of *δ*_1_ (the smaller value of v for *A/A*_0_ = 1 is more important than the smaller slope for A/A_0_ = 1). The relaxation time *τ* = 0.8 minutes is consistent with typical cortical turnover times [12]. This may imply that the relaxation of the transient tension originates mainly from the cortex. The large ratio *k_t_/k_ss_* = 63 indicates that the short-time elastic contribution of the transient tension dominates completely over the steady-state tension related to the same area increase. This is evident from the large slope of the red curve for small areas in Fig. 2b, compared to that of the dashed black curve. This ratio may suggest a change in the configuration of the membrane-cortex complex or its coupling to ion transporters during relaxation.

The volume loss during spreading was explained in Ref. [10] by transient tension alone. It is surprising, therefore, that our theory fits successfully the same data with a finite steady-state tension. In order to better understand this point, we focus on the data of Ref. [10] (green diamonds) at long times, when the predictions of the two theories are expected to differ most. Long times refer to larger areas in Fig. 2. The cells have not yet completed spreading and still have a finite spreading velocity. Both theories predict that there is a finite tension at such long times. The theory of Ref. [10] interprets it as a transient tension due to the spreading rate, which relaxes on the scale of ten minutes. The current theory interprets it as a steady-state elastic tension (dashed curve in Fig. 2), accompanied by a transient tension that relaxes quickly on the scale of one minute. Note that the final dimensionless volume in Fig. 2b is about 10% smaller than the initial one, while the final volume in Ref. [10] is only 3% smaller than the original one. This is due to cellular growth that is incorporated in *v*, as is explained above.

## IV. DISCUSSION

In this paper, we formulated a physical theory that captures the two different modes of cellular volume loss observed in the cell-spreading experiments of Refs. [8–10]. We explain the substantial differences between the experiments by a single difference between the cells: the cell density. Swollen cells, with a smaller fraction of dried volume, are more susceptible to volume loss, explaining the large volume loss in [8, 9], compared with [10]. This argument can be verified by repeating the experiments of Ref. [10] on treated cells that are sufficiently swollen due to modified ion transport rates.

While the important roles of cell activity and ion transport in the volume loss were clearly highlighted in the experiments of Refs. [8–10], the effect of mechano-sensitive channels was only partially demonstrated. It was shown [10] that inhibition of mechano-sensitive calcium channels using gadolinium chloride results in a smaller volume decrease. Further investigation of the role of mechano-sensitive channels in the effect is required.

Our theory is based on the existence of a steady-state tension increase with respect to a reference tension in suspension. This assumption can be verified by measuring the tension of spreading cells (*e.g*., using tether pulling or a fluorescent membrane-tension probe [24]). Such a tension can be driven, for example, by changes in the values of endo- and exocytosis rates or in the surface tension of lipid reservoirs in the cells. In Ref. [10], it was assumed that tension homeostasis is maintained, *i.e*., that the steady-state tension achieves its reference value with no residual tension due to spreading (*k_ss_* = 0). The steady-state volume is expected to be conserved in this case, in the absence of growth. Growth then leads to larger cells with a positive correlation between cell volume and spread area, as was reported also in Ref. [25].

Tether-pulling experiments from Ref. [10] did not indicate a steady-state tension increase. This is understandable in light of our fitted value *k_t_/k_ss_* = 63 that suggests that the steady-state modulus is very small compared to the transient one. It is expected to be difficult to detect, especially given the statistical error in such measurements. Also, since synchronization and starvation may drive cells out of their homeostatic state, it is possible that k_*ss*_ is larger for the starved and synchronized cells of Refs. [8, 9], compared with the non-treated cells of Ref. [10].

The relation between cell synchronization and starvation to cell swelling can be investigated further via a more detailed microscopic model (see Secs. A and B in Supplementary Material). Such a model identifies the chemical-potential difference *δ* with (minus) the membrane potential. More polarized cells (with larger membrane potentials in absolute values) have smaller volumes. It is known that the membrane potential evolves during the cell cycle [26] and, namely, that cells are hyperpolarized during the S and G2 phases. This may explain why synchronized cells have larger volumes.

Furthermore, potassium channels are known to have an especially important role in regulating the cell cycle [26], both by modifying the membrane permeability and via specific permeation-independent mechanisms. An ion-specific model, where the tension dependence originates mostly from mechano-sensitive potassium channels (see Sec. C of Supplementary Material), relates cell swelling to a lower potassium permeability. Such a model also predicts a more negative value of *δ*_1_ for swollen cells, compared to non-swollen cells, in accordance with our fits to the experiments.

Cell starvation and synchronization may have important implications on both the steady-state surface tension and cell swelling. Such cell treatments may also modify the activity of different ion pumps and the composition of trapped osmolytes, including the ratio *V_m_/V_d_*. For example, the same mass of trapped molecules (e.g., amino acids) with the same value of *V_d_* can either be bound in large macromolecules or be mobile (larger contribution to N) [27]. This implies that the value of *V_m_/V_d_* may change due to cell starvation and synchronization that alter protein synthesis rates. In this work, we assumed the same value of *V_m_/V_d_* for all experiments. A theoretical study of this effect must first be based on future experiments that investigate cellular volume regulation under different conditions of the trapped molecules (e.g., under different pharmacological treatments that modify protein synthesis rates). Since cell starvation and synchronization are often used in experiments, it is important to understand better how they modify cellular volume regulation. Our predictions can serve as basis for such studies in the future.

## ACKNOWLEDGMENTS

We thank Matthieu Piel for helpful discussions. RMA acknowledges funding from Fondation pour la Recherche Medicale (FRM Postdoctoral Fellowship). AVS and PS are supported by grants from Agence Nationale de la Recherche (MOTICAVANR-17-CE13-0020-01) and INSERM Cancer grant 20CR110-00. SAS is grateful for a Weizmann-Curie grant, the Volkswagen Foundation and the historic generosity of the Perlman Family Foundation.

## SUPPLEMENTARY MATERIAL

### A. Ion-specific pump-leak model

The pump-leak model presented in the main text considers neutral trapped molecules and neutral osmolytes that exchange with the buffer. In fact, the trapped molecules are negatively charged, on average, and the additional osmolytes consist of several ionic species. Here, we present an ion-specific model that accounts for these finer details. The model relates the chemical-potential difference *δ* to the membrane potential and provides physical insight regrading mechano-sensitivity.

Our ion-specific model considers a fixed number *N_p_* of trapped molecules that have an average electric charge –*ze*, where –*e* is the electron charge. This charge induces an electric potential difference *ψ* across the cell membrane (Donnan potential). Within the Donnan framework [15, 16], the electric field in the cell is negligible, and the potential is considered to be homogeneous inside the cell. The membrane potential attracts cations and repels anions that exchange with the buffer. For the sake of simplicity, we consider only the three dominant ionic species in the cell and buffer: Na+, K+, and Cl-. Na and K diffuse passively through channels and are pumped actively via the NaK pump that exports three Na ions and imports two K ions [6]. Cl transport is mostly passive and its active pumping is neglected hereafter. More detailed models can be considered (see, e.g., Ref. [?]).

As explained in the main text, mechanical equilibrium with the buffer is well approximated by osmotic pressure balance. The latter is given by

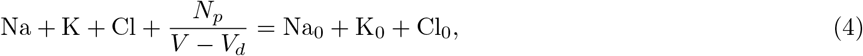

where Na, K, Cl are the cellular concentrations and Na_0_, K_0_, Cl_0_ are the buffer concentrations. Electroneutrality in the cell and in the buffer dictates

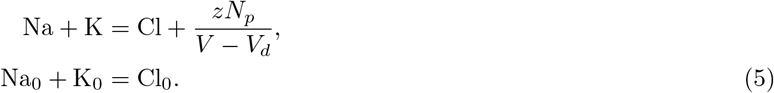

Inserting these relations in the osmotic-pressure equality yields

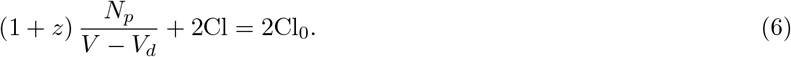

This equation can be understood via the coarse-grained model of the main text. The first term is the pressure of the trapped molecules and their counterions, which are overall neutral. The second term accounts for the additional ions in the cell, while the right-hand-side is the buffer pressure. Explicitly, the coarse-grained model is recovered by the substitutions (1 + *z*) *N_p_* → *N*, 2Cl → *n_c_*, and 2Cl_0_ → *n_b_*.

Since Cl ions are not pumped, the concentration of cellular Cl is related to the buffer concentration by chemical equilibrium,

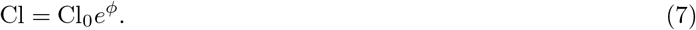

where *ϕ* = *eψ/k*_B_*T* is the dimensionless membrane potential. As was shown above, Cl is equivalent to the additional salt in the coarse-grained theory. Comparing to the relation *n_c_* = *n_c_* exp (–*δ*) shows that, within this model, *δ* = –*ϕ*. Indeed, substituting Eq. (7) in Eq. (6) leads to

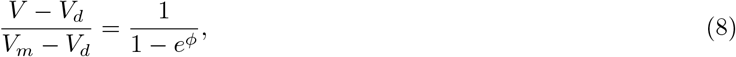

where *V_m_* = V_d_ + (1 + *z*) *N* / (2*Cl*_0_). Replacing *ϕ* → –*δ* restores Eq. (1) of the main text. Note that *V_m_* is the volume in the limit of an infinitely negative membrane potential, where no anions enter the cell (i.e., no added salt in the cell).

### B. Mechano-sensitivity from the microscopic model

The dimensionless membrane potential φ can be related to the ion flux and, by arguments of mechano-sensitivity, to tension. We take another linear combination of the pressure-equality and electroneutrality conditions [Eqs. (4) and (5), respectively],

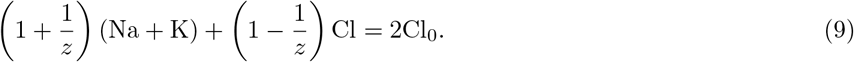

The cellular concentrations of Na and K can be related to their buffer concentrations from the the condition of zero net flux at steady state. In thermodynamic equilibrium, this condition reduces to chemical equilibrium, as is the case for the Cl ion [Eq. (7)]. For Na and K, on the other hand, chemical equilibrium is broken by the activity of the NaK pump. The electrochemical potential difference can be found within linear response as –3*r*/Λ_Na_ and 2*r*/Λ_K_ for Na and K, respectively, where *r* is the NaK-pump pumping rate per unit area of the cell surface and Λ_Na/K_ denotes the respective ion-channel conductance. This leads to

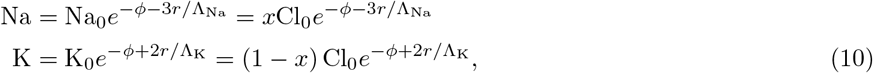

were *x* = Na_0_/Cl_0_ is the fraction of Na ions in the buffer, while 1 – *x* is the fraction of K ions.

Inserting Eq. (10) in Eq. (9) yields a relation between the membrane potential to the NaK pumping rate and the ion-channel conductance,

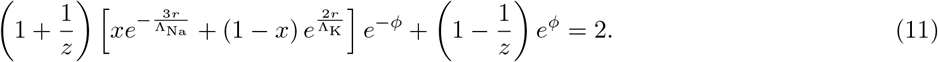

Generally, each of *r*, Λ_Na_, and Λ_K_ can be mechano-sensitive. The derivative of the potential with respect to the surface tension *dϕ/dγ* can be carried out in this general case [10] and it is expected to be negative under reasonable physical conditions.

Next, we relate the value of *ϕ* (that corresponds to –*δ* in the main text) to mechano-sensitive ion transporters. For simplicity, we consider monovalent trapped molecules *z* =1. We further neglect the cellular concentration of Na compared to K. This estimation is reasonable under standard physical conditions [6] and we consider that it holds also during spreading. In other words, we assume that *r*Λ_Na_ does not change much due to a tension increase. In this limit, Eq. (11) reduces to

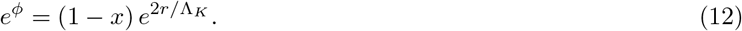

The potential is thus more negative for smaller *r*/Λ_K_ ratios. As mechano-sensitivity is attributed to channels more than to pumps, We consider only Λ_K_ to have a mechano-sensitive contribution. In this limit, the volume loss during spreading is due to an increased ion flux of K ions out of the cell through mechano-sensitive channels.

As a model for Λ_K_, we consider separate contributions from non-mechano-sensitive channels 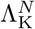 and mechano-sensitive channels 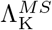,

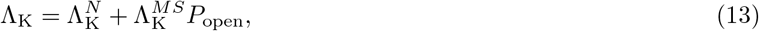

where *P_open_* is the probability for an open mechano-sensitive channel. We calculate this probability from a two-state model. Consider a mechano-sensitive potassium channel with two possible states, open and closed, and energies *ϵ*_open_ and *ϵ*_closed_, respectively. We associate the oppening of the transporter with a charecteristic membrane area *a*. The closed state, therefore, requires an additional energy *γa* to close the membrane area, given the surface tension *γ*.

The partition function of the mechano-sensitive channel is

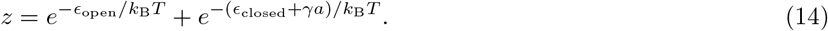

The probability for an open transporter is thus

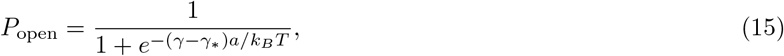

where *γ*_*_ = (*ϵ*_open_ – *ϵ*_closed_) /*a*. The tension *γ*_*_ is the inflection point of *P*_open_, and plays the same role as the Fermi energy in the Fermi-Dirac distribution. Close to *γ*\*, the open and closed states are almost equally probable, while applied tension favors the open state. Inserting Eqs. (15) and (13) for Λ_K_ in Eq. (12) for the membrane potential yields the relation *ϕ*(*γ*).

We expand *ϕ*(*γ*) around the state in suspension with tension *γ*_0_. This provides microscopic expressions for *δ*_0_ and *δ*_1_ of the main text. First, we expand the conductance, Λ_K_ = Λ_0_ + Λ_1_ (*γ* – *γ*_0_)/*γ*_0_, where

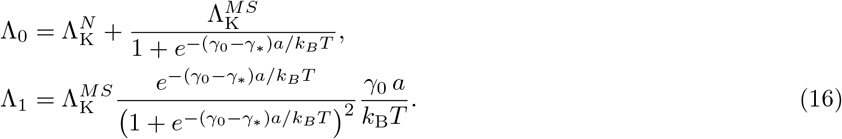

Next, expanding the potential as *ϕ* = *ϕ*_0_ + *ϕ*_1_ (*γ* – *γ*_0_) /*γ*_0_ yields

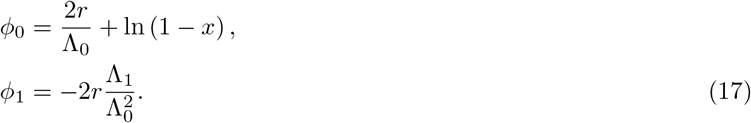

The potential in suspension is negative due to ln (1 – *x*), where 1 – *x* = K_0_/Cl_0_ is the ratio of the buffer concentrations of K and Cl. The result for *ϕ*_1_ shows that the potential becomes more negative as tension increases. This corresponds to *δ*_1_ > 0, as considered in the main text.

### C. Possible microscopic origin of swollen cells

Within this ion-specific model, swollen cells have a larger (less negative) membrane potential. According to Eq. (11), the potential is related to the Na and K conductance Λ_Na_ and Λ_K_, the active NaK pumping rate *r*, and the average valency of trapped molecules *z*. Changes in any of these parameters can lead to cell swelling.

We focus on our simple limit, where *z* =1 and where Λ_K_ is the main physical quantity that changes during spreading due to mechano-sensitivity. The potential in suspension is then given by Eqs. (16) and (18). Larger potentials correspond to either higher pumping rates or lower K conductance. Out of the two, lower K conductance seems more reasonable for starved, synchronized cells, as were used in Refs. [8, 9]. Explicitly, we consider the case where the conductance 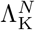 and 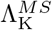 are lowered by a prefactor 0 < *α* < 1 that can originate, for example, from a smaller number of channels. In this case, both *ϕ*_0_ and *ϕ*_1_ are expected to change. We denote the potential change due to swelling as Δ*ϕ*, and find that

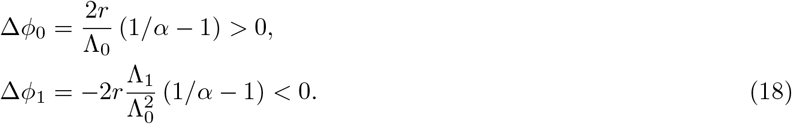

Here we have neglected changes in the potential due to a possible larger surface tension in the swollen state.

Equation (18) suggests that cells with a smaller number of channels have a smaller volume in suspension (less negative *ϕ*_0_), and are more susceptible to tension changes (more negative *ϕ*_1_). This augmentation of mechano-sensitivity refers to different values of *δ*_1_ in the main text and is complementary to the exponential augmentation due to different *δ*_0_ values.

## References

[1] C. Cadart, L. Venkova, P. Recho, M. C. Lagomarsino, and M. Piel, Nature Physics 15, 993 (2019).

[2] M. B. Ginzberg, R. Kafri, and M. Kirschner, Science 348, 1245075 (2015).

[3] E. K. Hoffmann, I. H. Lambert, and S. F. Pedersen, Physiological reviews 89, 193 (2009).

[4] D. Tosteson and J. Hoffman, The Journal of general physiology 44, 169 (1960).

[5] A. R. Kay, Frontiers in cell and developmental biology 5, 41 (2017).

[6] R. Milo and R. Phillips, Cell biology by the numbers (Garland Science, 2015).

[7] H. Jiang and S. X. Sun, Biophysical journal 105, 609 (2013).

[8] M. Guo, A. F. Pegoraro, A. Mao, E. H. Zhou, P. R. Arany, Y. Han, D. T. Burnette, M. H. Jensen, K. E. Kasza, J. R. Moore, F. C. Mackintosh, J. J. Fredberg, D. J. Mooney, J. Lippincott-Schwartz, and D. A. Weitz, Proceedings of the National Academy of Sciences of the United States of America 114, E8618 (2017).

[9] K. Xie, Y. Yang, and H. Jiang, Biophysical Journal 114, 675 (2018).

[10] L. Venkova, A. S. Vishen, S. Lembo, N. Srivastava, B. Duchamp, A. Ruppel, A. Williart, S. Vassilopoulos, A. Deslys, J.-M. G. Arcos, et al., eLife 11, e72381 (2022).

[11] R. M. Adar and S. A. Safran, Proceedings of the National Academy of Sciences 117, 5604 (2020).

[12] P. Chugh and E. K. Paluch, Journal of Cell Science 131(2018), 10.1242/jcs.186254.

[13] S. A. Safran, Statistical thermodynamics of surfaces, interfaces, and membranes (CRC Press, 2018).

[14] T. Markovich, D. Andelman, and R. Podgornik, Handbook of Lipid Membranes: Molecular, Functional, and Materials Aspects, 99 (2021).

[15] F. G. Donnan, Chemical reviews 1, 73 (1924).

[16] J. T. Overbeek, Prog Biophys Biophys Chem 6, 57 (1956).

[17] R. Phillips, J. Kondev, J. Theriot, et al., Physical biology of the cell (Garland Science, 2009).

[18] L. L. Norman, J. Bruges, K. Sengupta, P. Sens, and H. Aranda-Espinoza, Biophysical Journal 99, 1726 (2010).

[19] M. Doi, Soft matter physics (Oxford University Press, 2013).

[20] P. Kelly, Mechanics lecture notes: An introduction to solid mechanics (2022).

[21] A. R. Bausch, W. Moller, and E. Sackmann, Biophysical Journal 76, 573 (1999).

[22] E. Ponder and G. Saslow, The Journal of physiology 70, 18 (1930).

[23] E. Zhou, X. Trepat, C. Park, G. Lenormand, M. Oliver, S. Mijailovich, C. Hardin, D. Weitz, J. Butler, and J. Fredberg, Proceedings of the National Academy of Sciences 106, 10632 (2009).

[24] C. Roffay, G. Molinard, K. Kim, M. Urbanska, V. Andrade, V. Barbarasa, P. Nowak, V. Mercier, J. García-Calvo, S. Matile, et al., Proceedings of the National Academy of Sciences 118(2021).

[25] N. Perez Gonzalez, J. Tao, N. D. Rochman, D. Vig, E. Chiu, D. Wirtz, and S. X. Sun, Molecular biology of the cell 29 (2018).

[26] D. Urrego, A. P. Tomczak, F. Zahed, W. Stühmer, and L. A. Pardo, Philosophical Transactions of the Royal Society B: Biological Sciences 369, 20130094 (2014).

[27] R. Rollin, J.-F. Joanny, and P. Sens, bioRxiv (2022), 10.1101/2022.08.01.502021, https://www.biorxiv.org/content/early/2022/08/03/2022.08.01.502021.full.pdf.

